# Flatworm transcriptomes reveal widespread parasitism by histophagous ciliates

**DOI:** 10.1101/2023.09.17.558123

**Authors:** M. Ryan Woodcock, Kaleigh Powers, Kirsten Snead, Jason Pellettieri

**Affiliations:** Department of Biology, Keene State College, Keene, NH, USA; Department of Science, Mathematics & Technology, Medaille University, Buffalo, NY, USA; Department of Natural Science, Trocaire College, Buffalo, NY, USA; Ira A. Fulton Schools of Engineering, Arizona State University, Tempe, AZ, USA

## Abstract

Unicellular ciliates like *Tetrahymena* are best known as free-living bacteriovores, but many species are facultative or obligate parasites. These ‘histophages’ feed on the tissues of hosts ranging from planarian flatworms to commercially important fish and the larvae of imperiled freshwater mussels. Here, we developed a novel bioinformatics pipeline incorporating the nonstandard ciliate genetic code and used it to search for Ciliophora sequences in 34 publicly available Platyhelminthes EST libraries. From 2,615,036 screened ESTs, we identified nearly 6,000 high-confidence ciliate transcripts, supporting parasitism of seven additional flatworm species. We also cultured and identified *Tetrahymena* from nine terrestrial and freshwater planarians, including invasive earthworm predators from the genus *Bipalium* and the widely studied regeneration models *Dugesia japonica* and *Schmidtea mediterranea*. A cophylogenetic reconstruction provides strong evidence for coevolution of histophagous Ciliophora with their Platyhelminthes hosts. We further report the antiprotozoal aminoglycoside paromomycin expels *Tetrahymena* from *S. mediterranea*, providing new opportunities to investigate the effects of this relationship on planarian biology. Together, our findings raise the possibility that invasive flatworms constitute a novel dispersal mechanism for *Tetrahymena* parasites and position the Platyhelminthes as an ideal model phylum for studying the ecology and evolution of histophagous ciliates.

## INTRODUCTION

*Tetrahymena* is familiar to many biologists as a genus of pond-dwelling protozoans in which the Nobel Prize-winning discoveries of ribozymes and telomerase were made (Greider and Blackburn, 1985; Kruger et al., 1982). Despite sharing common morphological features, these unicellular ciliates exhibit a remarkable and underappreciated degree of biodiversity. The recent application of DNA barcoding approaches has distinguished more than 80 species, and these probably represent only a small fraction of the total in existence today (Chantangsi et al., 2007; Doerder, 2019). While many inhabit temperate, freshwater ecosystems, they have also been documented in the Arctic and Antarctic, moss and soil samples, carnivorous plants, and parasitizing a wide variety of invertebrate and vertebrate hosts (Brandl et al., 2005; Doerder, 2019; Elliott et al., 1962; Petz et al., 2007; Pitsch et al., 2017; Stout, 1954; Strüder-Kypke et al., 2001).

Traditionally, *Tetrahymena* were categorized into three major groups, or complexes, based upon their life history traits – the predominantly bacterivorous *pyriformis* complex, the predatory *patula* complex, members of which consume other ciliates, and the parasitic *rostrata* complex, consisting of histophagous forms that feed on the tissues of their hosts (Corliss, 1970). Molecular phylogenies do not resolve these complexes into discrete clades, however, and instead support parallel evolution of the above traits within the *Tetrahymena* genus (Doerder, 2019; Lynn et al., 2018; Rataj and Vďačný, 2020; Strüder-Kypke et al., 2001). While recent reconstructions have led to different conclusions about whether bacterivory or histophagy was the ancestral life history, transitions between the two almost certainly occurred repeatedly during diversification of this lineage (Lynn et al., 2018; Rataj and Vďačný, 2020; Strüder-Kypke et al., 2001).

Histophagous *Tetrahymena* prey upon a broad range of invertebrates, including midges (*T. chironomi*) (Corliss, 1960), mosquitoes (*T. empidokyrea*) (Jerome et al., 1996), black flies (*T. dimorpha, T. rotunda*) (Batson and Beale, 1983; Lynn et al., 1981), alderflies (*T. sialidos*) (Batson and Rees, 1985), slugs (*T. limacis, T. rostrata*) (Brooks, 1968), the glochidia larvae of freshwater mussels (*T. glochidiophila*) (Lynn et al., 2018), and the planarian flatworms *Crenobia alpina, Dendrocoelum lacteum, Dugesia gonocephala, Girardia tigrina, Phagocata vitta*, and *Polycelis felina* (*T. acanthophora, T. corlissi, T. dugesiae, T. nigricans, T. pyriformis, T. scolopax*) (Armitage and Young, 1990; Rataj and Vd’ačný, 2019; Rataj and Vďačný, 2020; Wright, 1981).

Vertebrate infections have been reported in fish (*T. corlissi, T*. sp.) (Astrofsky et al., 2002; Ferguson et al., 1987; Hoffman et al., 1975; Leibowitz and Zilberg, 2009), frogs (*T*. sp.) (Holz et al., 2015; Shaw et al., 2011), salamanders (*T*. sp.) (Shumway, 1940), chicks (*T. pyriformis*) (Knight and McDougle, 1944), and even a dog (*T. farleyi*) (Lynn et al., 2000). Although there may be some commensal relationships (Lynn et al., 2000; Pitsch et al., 2017), most of the above associations have been interpreted as cases of facultative or constitutive parasitism (in the former, parasites can also propagate independently of their host) (Doerder, 2019).

*Tetrahymena* infections are often associated with tissue lesions and mortality, and a causal link is supported by laboratory studies. For example, the larvae of freshwater mussels experienced a rapid decline in viability when exposed to cultured *T. glochidiophila* (Prosser et al., 2018), as did ornamental fish exposed to *T*. sp. parasites responsible for a condition known as ‘guppy killer disease’ (Sharon et al., 2014). Given the threatened or endangered status of many unionid mussel species and the commercial importance of the ornamental fish trade, these results underscore the potential for histophagous *Tetrahymena* and related ciliates to have damaging ecological and economic impacts.

We now report a bioinformatic analysis of platyhelminth EST databases substantially expanding the known associations between ciliates and flatworms, as well as the isolation and identification of *Tetrahymena* from nine wild-caught and laboratory-maintained planarian populations. Some of the infected species are invasive, raising the possibility they contribute to the geographic dispersal of *Tetrahymena* parasites. Together with other recent studies (Rataj and Vd’ačný, 2021, 2019; Rataj and Vďačný, 2020), our results support frequent histophagy in the Ciliophora and establish the Platyhelminthes as an ideal model phylum for investigating coevolution between parasitic ciliates and their hosts.

## RESULTS

### EST libraries support widespread association of flatworms and ciliates

Motivated by the observation of apparent *Tetrahymena* transcripts in a planarian RNA-seq dataset (He et al., 2017), we developed a bioinformatic pipeline to search for ciliate sequences in publicly available Platyhelminthes EST libraries (Figure 1A; Materials and Methods). In the first step, 2,615,036 input ESTs from 32 flatworm species (Table S1) were translated using the ciliate genetic code, in which codon reassignment results in translation of standard UAA and UAG stop codons as glutamine (Horowitz and Gorovsky, 1985; Preer et al., 1985). Predicted protein sequences were then assigned preliminary taxonomic classifications using GhostKOALA, an automatic annotation and KEGG mapping server developed for rapid characterization of metagenomes (Kanehisa et al., 2016). This revealed 710,805 candidate ciliate ESTs, which were next subjected to a pair of more stringent selection criteria. First, they had to exhibit greater similarity to Ciliophora than to Platyhelminthes sequences in unidirectional NCBI BLASTP searches (queries were translated in the ciliate code for Ciliophora BLASTs and the standard code for Platyhelminthes BLASTs). Second, they had to be identified as orthologs of annotated *T. thermophila* genes, using the orthologous clustering tool OrthoVenn2 (Xu et al., 2019). Application of these filters produced a set of 6,392 predicted ciliate ESTs, representing 5,957 unique *T. thermophila* orthologs (Table S2).

**Figure 1:**
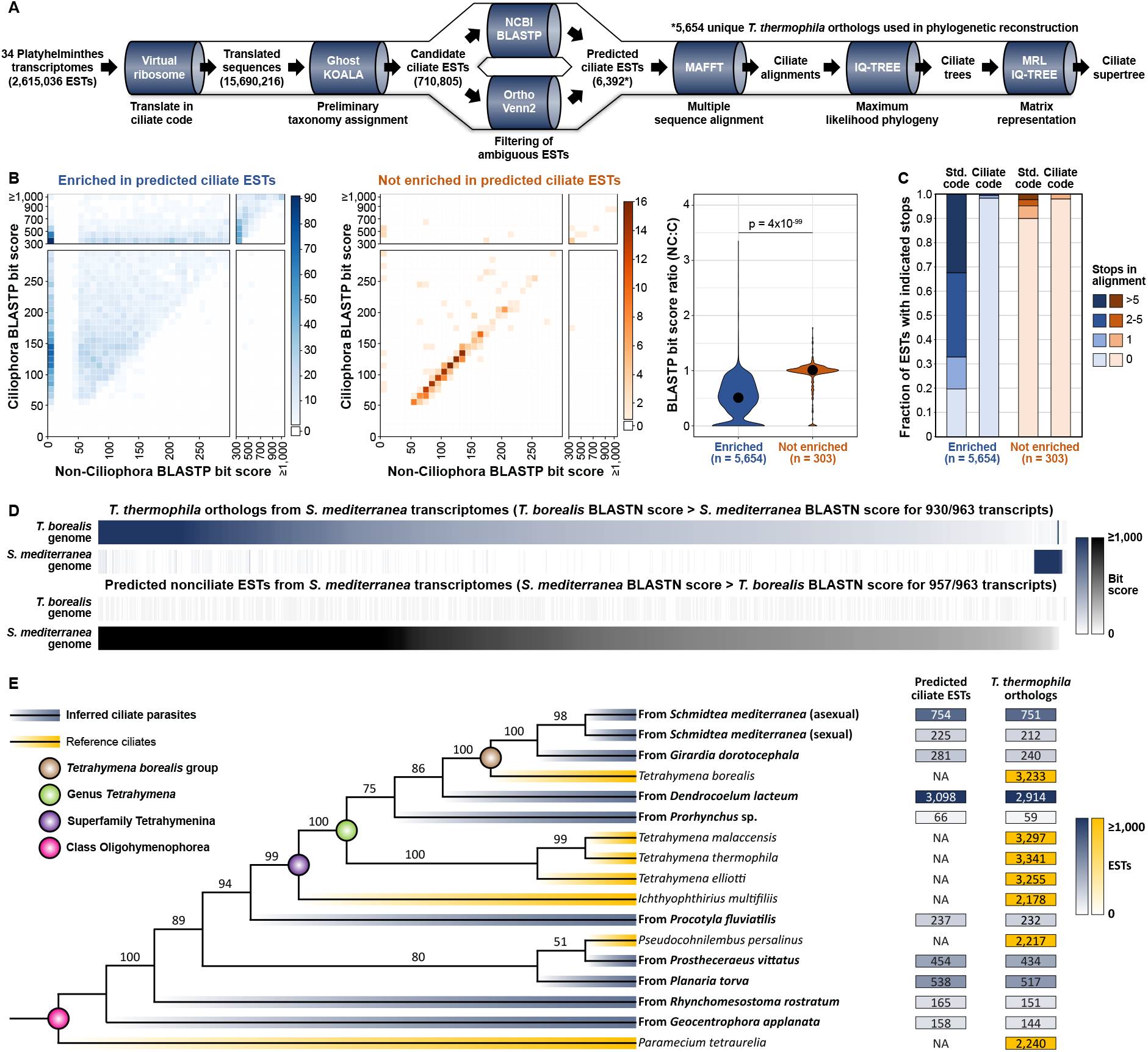
Identification and phylogenetic analysis of ciliate ESTs in flatworm transcriptomes. **A.** Schematic illustration of sequence analysis pipeline (see Materials and Methods for details). All candidate ciliate ESTs identified by GhostKOALA were evaluated by BLASTP and OrthoVenn2 in parallel; only those meeting criteria for both were designated as predicted ciliate ESTs. Unique *T. thermophila* orthologs from this dataset were used for phylogenetic reconstruction when the number of predicted ciliate ESTs in a given transcriptome exceeded the calculated false-positive threshold (see Table S3). **B**. 2D histograms and violin plots depicting BLASTP results for *T. thermophila* orthologs recovered from transcriptomes enriched (blue) or not enriched (orange) in predicted ciliate ESTs. Query sequences were translated in the ciliate code for BLASTs against the Ciliophora database and the standard code for BLASTs against the non-Ciliophora database. Color scales in 2D histograms denote number of transcripts. Violin plots reflect the distribution of non-Ciliophora (NC):Ciliophora (C) bit score ratios, with circles indicating medians. P-value is for two-tailed T-test. **C**. Stacked bar chart showing the fraction of *T. thermophila* orthologs with the indicated numbers of stop codons falling within the aligned region of the top Ciliophora BLASTP hit. **D**. Heat maps depicting bit scores for BLASTN alignments of the indicated transcripts against the *S. mediterranea* and *T. borealis* genomes. **E**. Maximum likelihood supertree generated from predicted ciliate ESTs. *Tetrahymena thermophila* open reading frames and orthologs from inferred ciliate parasites (blue) or reference species (yellow) were translated in the ciliate code and used to construct multiple sequence alignments in MAFFT. Individual alignments were then concatenated and used to infer a maximum likelihood supertree from molecular phylogenies in IQ-TREE. Nonparametric bootstrap frequencies were calculated from 10,000 replications. Heat-mapped data at right indicate number of transcripts. BLAST scores and stop codon numbers for **B**,**C**,**D** are included in source data files.

To estimate the false-positive rate for our approach, we also ran EST libraries from five unrelated eukaryotes, selected partly on the basis of being highly unlikely ciliate hosts, through our pipeline (Table S3; Supplementary Note 1). Relative to these transcriptomes, which yielded a calculated false-positive rate of 0.08%, 10 of the 34 screened Platyhelminthes libraries showed significant enrichment in predicted ciliate ESTs (Table 1). The corresponding 5,654 unique *T. thermophila* orthologs exhibited multiple features consistent with being of ciliate origin, in addition to meeting all pipeline criteria. First, 97% produced higher bit scores for BLASTP alignments against Ciliophora sequences than against all non-Ciliophora sequences in the NCBI nonredundant database (Figure 1B, blue). In contrast, the 303 *T. thermophila* orthologs from libraries not enriched in predicted ciliate ESTs (i.e., potential false positives) generally had similar Ciliophora and non-Ciliophora BLASTP scores, with 52% generating a better non-ciliate hit (Figure 1B, orange). Independent taxonomic classification using an algorithm that scores exact matches in protein sequences (Kaiju) (Menzel et al., 2016) yielded comparable results, with over 92% of classified ESTs from the 10 enriched transcriptomes assigned to the Ciliophora phylum (Materials and Methods; Figure S1). Second, 80% of the *T. thermophila* orthologs from the enriched dataset had one or more stop codon(s) within the aligned region of the top Ciliophora BLASTP hit when the query sequence was translated in the standard code, but this number fell to less than 2% when the query was translated in the ciliate code (Figure 1C, blue). Conversely, the large majority of potential false-positive sequences (90%) had no stops within the aligned region regardless of which code was used (Figure 1C, orange). Third, 930/963 (97%) of the *T. thermophila* orthologs recovered from the two screened *S. mediterranea* transcriptomes produced higher BLASTN bit scores when queried against a *Tetrahymena* (*T. borealis*) genome (Yang et al., 2019) than against the *S. mediterranea* genome (Grohme et al., 2018), whereas the opposite was true for a randomly selected set of control *S. mediterranea* transcripts excluded at the GhostKOALA step of the pipeline (Figure 1D). Finally, manual inspection revealed the presence of numerous ciliate-specific genes, such as granule lattice and tip proteins (Rahaman et al., 2009), within our list of pipeline-selected ESTs (Table S2). In conclusion, we consider the 5,654 unique *T. thermophila* orthologs from the 10 enriched Platyhelminthes transcriptomes to be high-confidence Ciliophora transcripts. While there may be bona fide ciliate sequences among those selected from the remaining 24 libraries (or those filtered during transcriptome assembly – see Supplementary Note 2), these also appear to include highly conserved genes that happened to produce marginally better Ciliophora than Platyhelminthes matches in BLASTP searches. Thus, only the enriched (high-confidence) dataset was subject to further analysis.

**Table 1:**
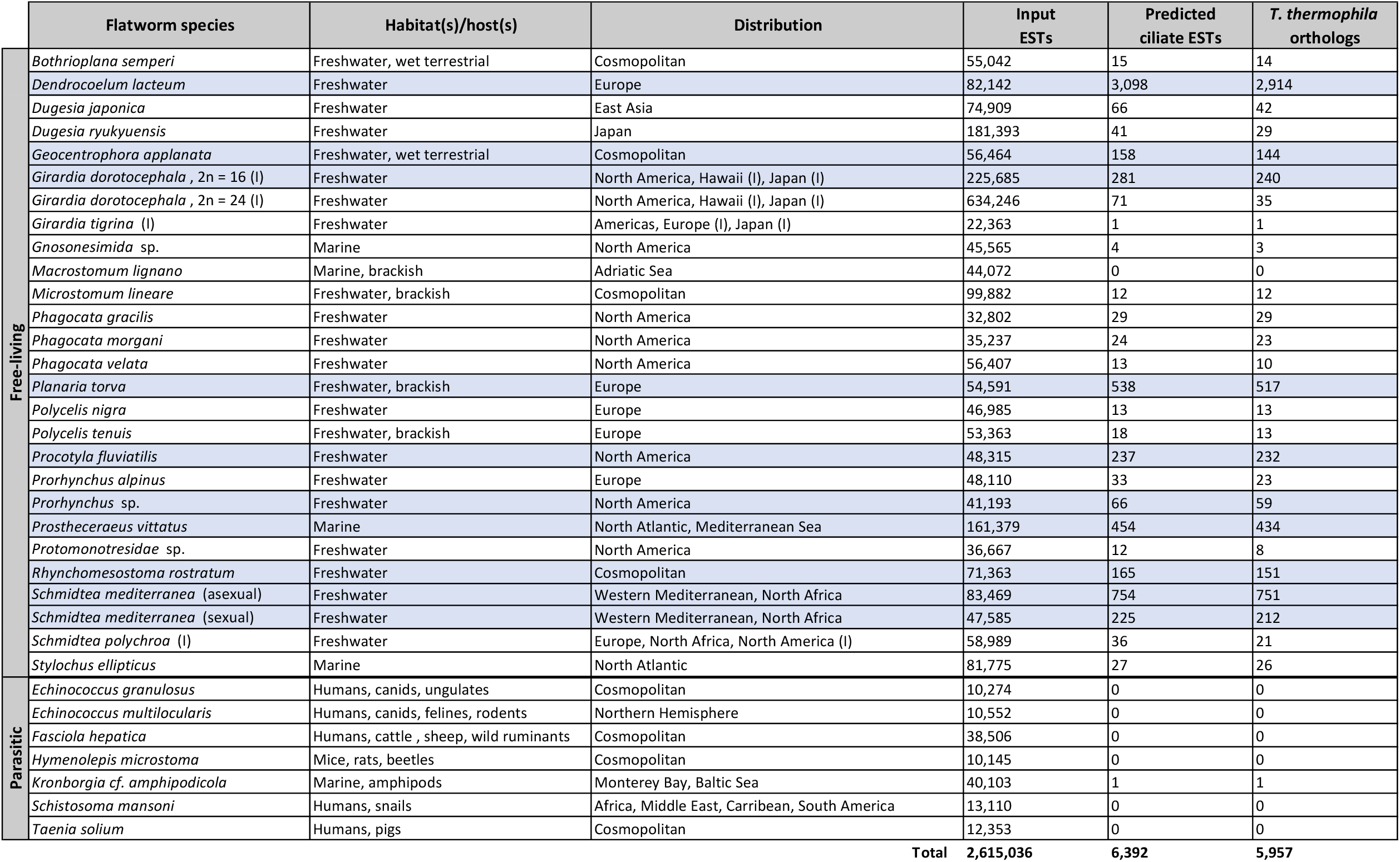
Identification of ciliate ESTs in flatworm transcriptomes. A total of 34 Platyhelminthes transcriptomes (see Table S1 for sources) were screened for ciliate ESTs using the bioinformatic pipeline outlined in Figure 1A. Invasive species and non-native distributions are marked by (I). Transcriptomes with a significantly higher number of predicted ciliate ESTs than expected false positives are shaded blue.

Our pipeline results did not allow for species identification; however, we were able to reconstruct a molecular phylogeny for the inferred ciliate parasites by comparing recovered ESTs to orthologous sequences from annotated reference species (Materials and Methods; Supplementary Note 3). Five branches of a maximum likelihood supertree generated by alignment of concatemerized translated sequences were assigned to the genus *Tetrahymena* with strong bootstrap support; the remainder were outgrouped with other oligohymenophoreans, including the fish pathogens *Ichthyophthirius multifiliis* and *Pseudocohnilembus persalinus* (Figure 1E). Kaiju classification of the high-confidence ciliate transcripts reinforced this overall organization (Figure S1). Together, the above results provide added evidence for prevalent infection of flatworms by Ciliophora parasites and establish a novel, bioinformatic approach for detecting these relationships (Discussion).

### Culture and identification of histophagous *Tetrahymena* infecting planarians

As a complement to our bioinformatic analysis, we isolated, cultured, and identified ciliates infecting a variety of wild-caught and laboratory-maintained planarian populations (Materials and Methods). Animals were thoroughly rinsed and dissected in water, after which any released ciliates were provided mealworms as a food source [commonly used for propagation of histophagous strains (Strüder-Kypke et al., 2001)]. DNA was extracted from cultures after 48 hours of growth for species identification by mitochondrial *cytochrome-c oxidase subunit 1* (*cox1*) barcoding (Chantangsi et al., 2007; Doerder, 2019). We identified *Tetrahymena* infecting an additional seven planarian species using this approach (Figure 2).

**Figure 2:**
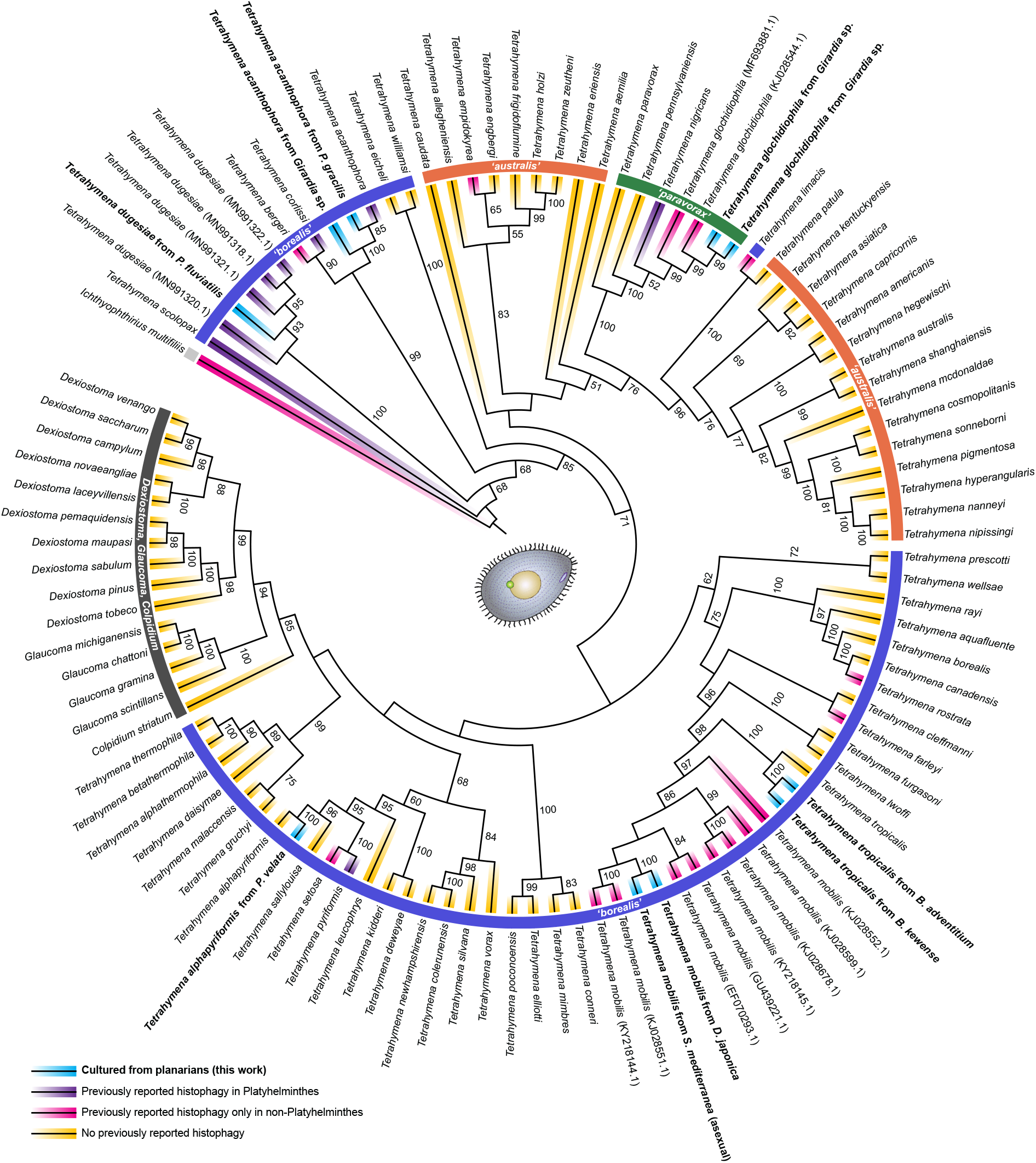
Phylogenetic analysis of *Tetrahymena* parasites cultured from planarians. Mitochondrial *cox1* DNA sequences from reference species and *Tetrahymena* isolates were aligned by MAFFT using RevTrans 2.0 to ensure alignment of orthologous codon positions. The resulting multiple sequence alignment was then used to infer a maximum likelihood phylogeny in IQ-TREE. Branch support analysis was conducted with 10,000 bootstrap replications (unmarked branches have bootstrap values of <50). Species identification was based on phylogenetic position and NCBI BLASTN hits. The two *T. glochidiophila* isolates were from two different animals of the same *Girardia cf. tigrina* population; the *T. acanthophora* isolate is from *Girardia cf. dorotocephala* (Materials and Methods). *cox1* sequences and accession numbers are included in the source data file.

Isolates were recovered from two of the three major *Tetrahymena* clades (‘borealis’ and ‘paravorax’), emphasizing the deep evolutionary relationship between the Platyhelminthes and Ciliophora (see below). Six of the new associations expand the known host range of previously documented histophages (*T. acanthophora, T. dugesiae, T. glochidiophila*, and *T. mobilis*), while the others imply at least some degree of histophagy among species commonly regarded as free-living bacteriovores (*T. alphapyriformis* and *T. tropicalis*). We were notably able to culture *Tetrahymena* from three flatworm species that were not enriched in predicted ciliate ESTs in our bioinformatic analysis (*D. japonica, P. gracilis*, and *P. velata*); this could reflect actual differences in the occurrence of ciliate infections among different populations or could be a byproduct of transcriptome filtering and assembly methods (Supplementary Note 2).

### Coevolution of ciliates and their flatworm hosts

Our results nearly double the number of flatworm species with documented ciliate parasites, establishing the Platyhelminthes as a heavily infected host phylum. To further explore evolutionary relationship between the Ciliophora and Platyhelminthes, we conducted a cophylogenetic analysis using Jane, an event-based reconstruction tool that finds optimal solutions for reconciling paired trees (Conow et al., 2010) (Materials and Methods). A total of 29 flatworms and 33 ciliates, representing all determined associations (30 previously identified and another 19 from the current study), were included in our analysis (Figure 3). There were 11 flatworms and three ciliates with multiple associations. While these numbers are somewhat complicated by the inclusion of results from studies conducted prior to the introduction of molecular phylogenetics (with consequent potential for misidentification of ciliate species), the data provide unambiguous support for host-parasite coevolution. Specifically, we observed significantly greater congruence between the Platyhelminthes and Ciliophora phylogenies than would be expected by chance, using either of two different randomization approaches (random tip mapping or random parasite trees) and any of seven different applied cost schemes (Table S4; Figure S2). Both vertical and horizontal transmission of ciliates were evident during evolution of their flatworm hosts, with at least one cospeciation event and multiple host switches predicted for every cost scheme. Thus, some or all of the host-parasite pairs appear to represent tight, long-term associations, but phylogenetic host specificity within the Platyhelminthes must be low enough to have allowed for frequent parasite transmission from one species to another (Discussion).

**Figure 3:**
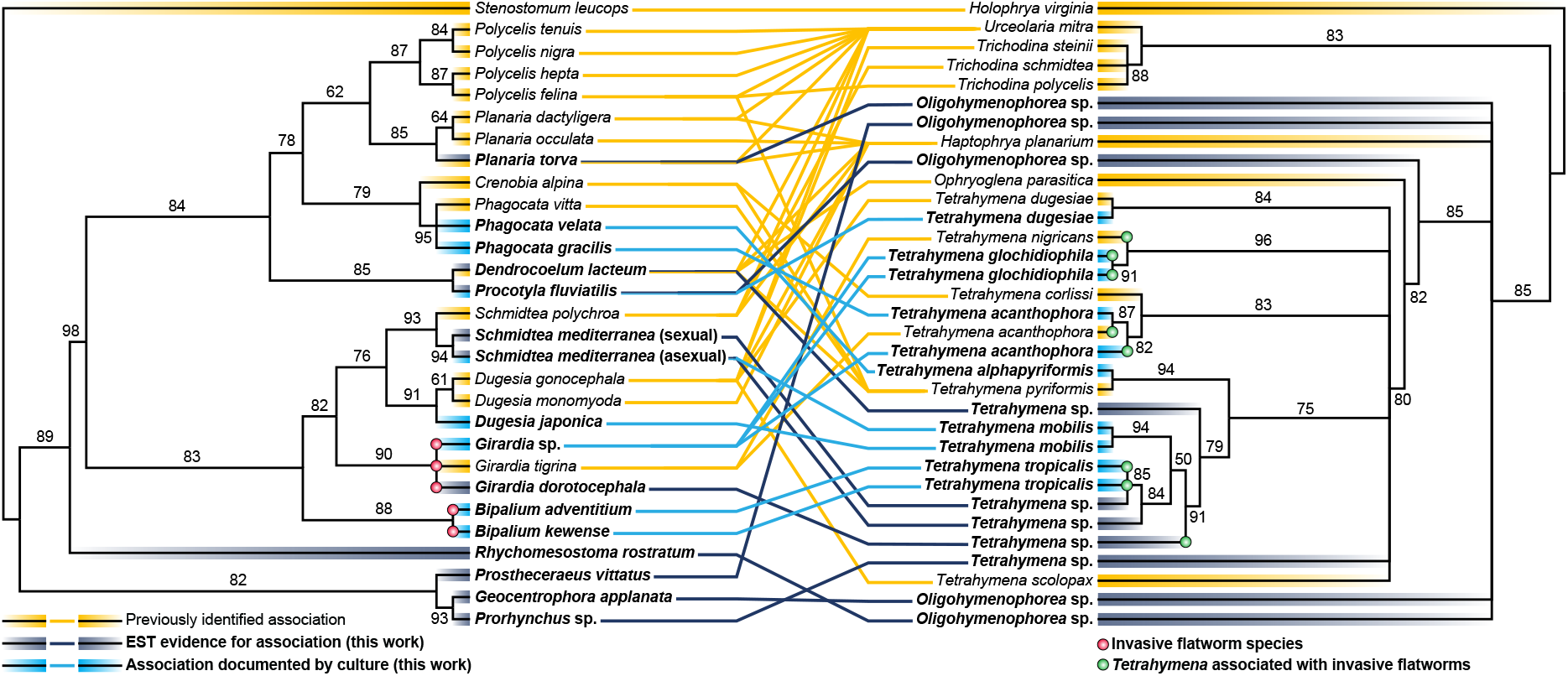
Cophylogeny of ciliate parasites and their flatworm hosts. Maximum likelihood supertrees for the Platyhelminthes and Ciliophora were generated from multiple, partitioned datasets using MAFFT and IQ-TREE (see Materials and Methods for data sources). Branch support analysis was conducted with 10,000 bootstrap replications. TreeMap was then used to connect each supertree and illustrate host-parasite associations (Platyhelminthes taxa not having known ciliate associations were pruned prior to this step).

### Histophagous *Tetrahymena* infecting planarians are facultative parasites

Some histophagous ciliates, such as *Ichthyophthirius multifiliis*, are obligate parasites that cannot survive for extended periods of time outside of their hosts (Dickerson and Findly, 2014). Our ability to obtain enough DNA for *cox1* barcoding from mealworm cultures (Figure 4A) suggests the *Tetrahymena* we isolated from planarians do not fall into that category. To confirm this hypothesis, we directly assessed the capacity of several different strains to proliferate in a free-living state. All tested isolates underwent exponential growth over the first three days in axenic culture, though some showed a plateau thereafter (Figure 4B). We conclude that the cultured *Tetrahymena*, including the two *T. tropicalis* strains recovered from invasive *Bipalium* species, are facultative parasites capable of at least limited growth independent of their hosts. This has major implications for potential dispersal mechanisms (Discussion).

**Figure 4:**
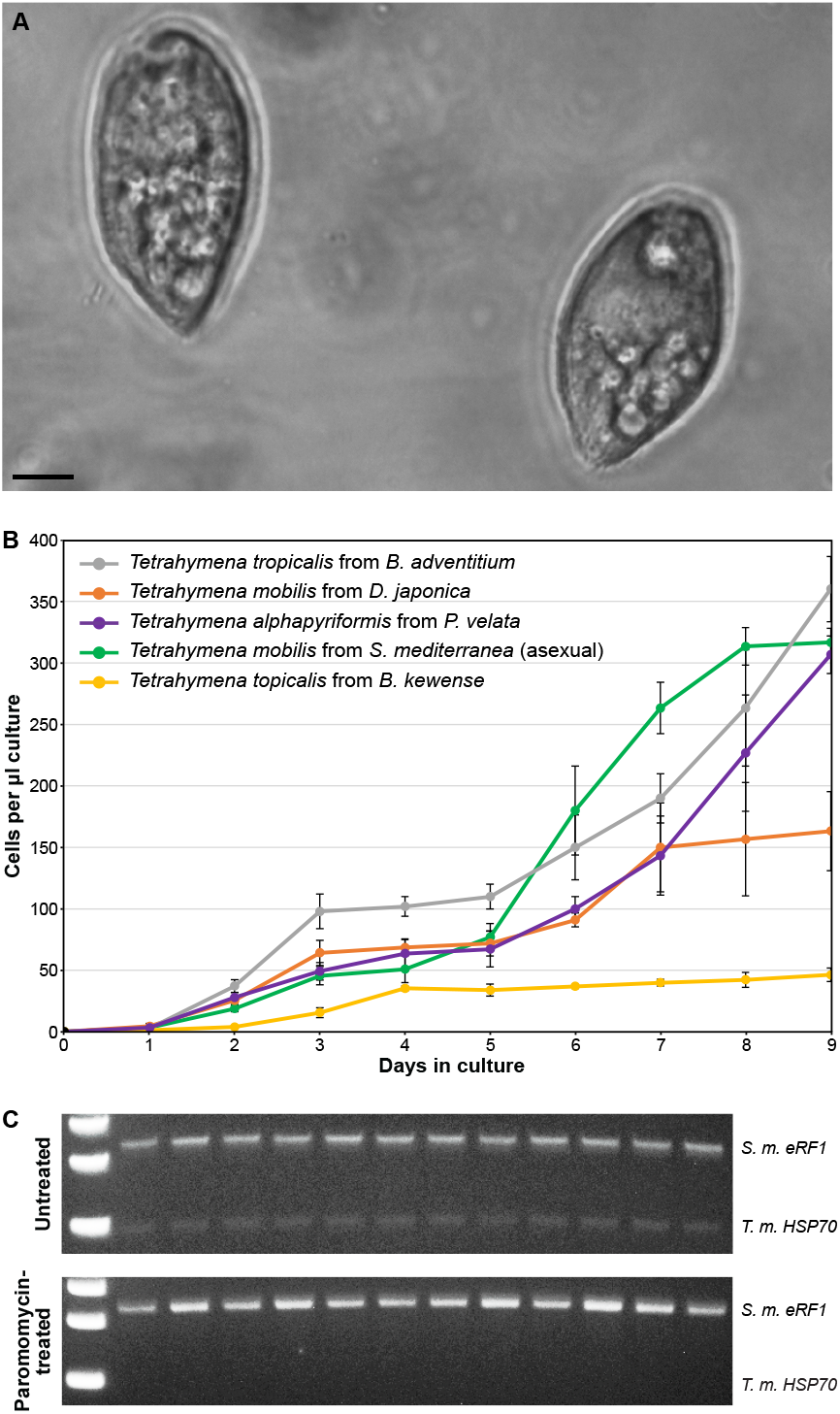
*Tetrahymena* isolates from planarians are facultative parasites. **A.** *Tetrahymena mobilis* cultured from *S. mediterranea* with mealworm as a food source. Scale bar = 10 µm. **B**. Growth curves for *Tetrahymena* isolates in Neff medium. Each data point is an average of three biological replicates. Error bars show standard deviation. **C**. PCR assay for *T. mobilis* (*T. m*.) infection in *S. mediterranea* (*S. m*.). Each lane to the right of the ladder shows PCR products (or lack thereof) amplified from total DNA templates obtained from individual animals maintained under standard laboratory conditions (top) or treated with paromomycin for a total of 28 days (bottom). PCR results were confirmed by mealworm culture (10 animals per condition; 100% positive for controls and 100% negative for paromomycin-treated). Cell counts and original gel images for **B**,**C** are included in source data files.

Failure to maintain optimal culture conditions can lead to the development of superficial tissue lesions in laboratory populations of *S. mediterranea*, and this often correlates with the appearance of large numbers of ciliates in the water (Merryman et al., 2018). To begin investigating possible adverse impacts of *Tetrahymena* infection, we sought to develop an effective antiprotozoal treatment. Vigorously rinsing animals in fresh water had no obvious influence on our ability to culture *T. mobilis*, even when this was repeated periodically over several days. We therefore tested the effects of maintaining planarians in the aminoglycoside antibiotic paromomycin, an antiprotozoal drug that has been shown to inhibit the growth of *T. thermophila* in culture and is used to treat the infectious protozoan disease leishmaniasis (Palmer and Wilhelm, 1978; Pokharel et al., 2021). One month of continuous paromomycin exposure abolished our ability to detect *T. mobilis* using a PCR-based assay (Figure 4C), without causing any evident host toxicity. Treated animals did eventually test positive again after treatment was stopped; it is unclear whether this was due to renewed growth of ciliates that survived drug exposure or re-infection. In either case, paromomycin represents a useful new tool that sets the stage for future studies addressing the impacts of *Tetrahymena* on various aspects of planarian biology.

## DISCUSSION

Prior to the discovery of their different mating type systems and resulting reproductive isolation, many morphologically similar *Tetrahymena* species were grouped together as *T. pyriformis* (Nanney and Simon, 2000). Today, it is clear these (and other) unicellular ciliates exhibit considerable ecological and evolutionary diversity. For example, while the majority of described species have only been observed in a free-living state, there are also a growing number of recognized endosymbionts that live in association with other organisms (Doerder, 2019). The bioinformatic pipeline we report here stands to accelerate the identification of new ciliate parasites and the animals they infect. We show that Ciliophora sequences can be readily recovered from the transcriptomes of their hosts, in part by translating ESTs in the nonstandard ciliate genetic code (Figure 1A-D). These sequences can subsequently be used for phylogenetic and co-phylogenetic reconstruction (Figures 1E and 3), though it is important to acknowledge that the potential advantages of multigene analyses are to some extent offset by the complications that could arise from a simultaneous infection by multiple ciliate species (Supplementary Note 3). Screening of all 34 Platyhelminthes EST libraries was carried out on public sites running specific bioinformatic tools and the open-source Galaxy server (The Galaxy Community, 2022), emphasizing the accessibility of our approach.

The 19 new ciliate-flatworm associations we established in the present study provide added evidence for coevolution between these phyla. Event-based models support between 13 and 25 host switches and one to 17 cospeciations (Table S4). Host switches are consistent with low phylogenetic host specificity among ciliates infecting the Platyhelminthes (Rataj and Vd’ačný, 2021; Rataj and Vďačný, 2020). Cospeciation indicates at least some interactions should be interpreted as long-term evolutionary relationships, rather than chance, temporary infections. *Tetrahymena mobilis*, for example, has persisted in laboratory populations of *S. mediterranea* for decades, cannot be eliminated by repeated washes, and was detectable in every animal we screened in our colony (Figure 4C).

Although we cannot fully exclude commensal or mutualistic relationships (see below), available evidence suggests the ciliate-flatworm associations that we and others have identified are cases of facultative parasitism. Most of the *Tetrahymena* isolates we recovered are known facultative histophages [e.g., *T. glochidiophila* (Prosser et al., 2018)], and all could be maintained in culture with mealworms as a food source. This is consistent with previous reports that *Tetrahymena* can feed directly on planarian tissues (Rataj and Vd’ačný, 2019; Rataj and Vďačný, 2020; Wright, 1981). The two isolates not previously identified as parasites, *T. alphapyriformis* and *T. tropicalis*, do have close histophagous relatives, including *T. mobilis*, which we now show infects both *D. japonica* and *S. mediterranea* (Figure 2). Indeed, the fact that even the well-known bacteriovore *T. thermophila* is capable of consuming animal cells in vitro may indicate at least some capacity for facultative histophagy is broadly conserved (Pinheiro and Bols, 2014). Future studies are likely to further expand the range of ciliates known to feed on animal tissues, as well as the flatworms they are capable of infecting. One notable exception may concern platyhelminth parasites – we did not detect evidence for ciliate associations with any of the six endoparasitic flatworms we surveyed (Table 1), perhaps reflecting more limited opportunities for histophages to prey upon species that themselves reside within a host.

One longstanding question about ciliates concerns their mechanisms of dispersal. Some form cysts, which facilitate survival in harsh environments and redistribution to new locations partly through attachment to organisms such as mites (which employ a similar strategy for their own dispersal) (Bharti et al., 2020). How cystless ciliates are spread, particularly over long distances, is not as well understood (Doerder, 2019). The infection of invasive *Bipalium* and *Girardia* species (Figure 3) may be relevant to this issue. The former are terrestrial, ‘hammerhead’ planarians native to Asia but now observed worldwide as a result of human transport of plants and soil (Justine et al., 2019, 2018). *Girardia dorotocephala* and *G. tigrina* are freshwater planarians that have spread from the Americas to Europe, Hawaii, and Japan (Kanana and Riutort, 2019; Kawakatsu et al., 1984; Sluys et al., 2010) (Table 1). As these (and perhaps other) invasive flatworms colonize new ecosystems, they have the potential to bring their ciliate parasites with them. An analogous scenario has been proposed for *Trichodina mutabilis*, a ciliate pathogen of numerous fish that has attained a global distribution via transcontinental shipments of carp (Basson and Van As, 2006; Islas-Ortega and Aguilar-Aguilar, 2014).

The ability to infect diverse hosts (Figure 3), while also proliferating independently of them (Figure 4B), may exacerbate the impacts ciliate parasites have on any new ecosystems to which they are transported. For example, we now show that *T. glochidiophila* infects *Girardia cf. tigrina* (confirmed by isolation from two separate animals), in addition to freshwater mussel larvae (Lynn et al., 2018). With the established capacity to propagate in bacterized culture (Prosser et al., 2018), any *T. glochidiophila* dispersed by flatworms could in theory proliferate in a free-living state before going on to infect new molluscan hosts. Many unioid mussels are threatened or endangered keystone species in freshwater ecosystems (Haag and Williams, 2014), so any resulting decline in viability (Prosser et al., 2018) could have far-reaching consequences.

Further investigations of how ciliates impact flatworms, and vice versa, will benefit from the many experimental tools that have been developed for *S. mediterranea*, a widely studied model for research on stem cell biology and regeneration (Ivankovic et al., 2019). This now includes the use of paromomycin to reduce *T. mobilis* numbers below the limit of detection by PCR (Figure 4C). *Tetrahymena* have been isolated from the gastrovascular tissues of other planarians (Rataj and Vďačný, 2020), and distantly related ciliate endosymbionts play an important role in ruminant digestion (Ushida, 2018). Thus, it will be interesting to determine whether paromomycin-treated *S. mediterranea* show any changes in food consumption and/or growth, in addition to determining the typical number and distribution of *T. mobilis* cells within the planarian anatomy. Results from such studies will complement the analyses reported and referenced here to provide a better understanding of how ciliate parasites have coevolved to exploit the widespread association with their Platyhelminthes hosts.

## MATERIALS AND METHODS

### Identification of predicted ciliate ESTs in flatworm transcriptomes

#### VirtualRibosome

Input ESTs for the pipeline analysis (Table S1) were first translated in VirtualRibosome 2.0 (Wernersson, 2006) in all six reading frames using the ciliate nuclear genetic code (translation table 6). Selected sequences were also translated in the standard genetic code (translation table 1) for BLASTP searches (see below).

#### GhostKOALA

Initial taxonomy classifications were made by submitting ESTs translated by VirtualRibosome to GhostKOALA (Kanehisa et al., 2016) for KEGG orthology assignments. All available prokaryotic and eukaryotic sequences were included in the search database (Platyhelminthes and Ciliophora included in the KEGG species list were *Echinococcus granulosus, Opisthorchis viverrine, Schistosoma haematobium, Schistosoma mansoni, Paramecium tetraurelia*, and *Tetrahymena thermophila*). Input sequences scoring as most closely related to *Tetrahymena* or *Paramecium* genes were classified as candidate ciliate ESTs.

#### NCBI BLASTP

Candidate ciliate ESTs were aligned against NCBI protein databases in BLASTP searches, with queries against Platyhelminthes sequences (Taxonomy ID: 6157) translated in the standard code and queries against Ciliophora sequences (Taxonomy ID: 5878) translated in the ciliate code. Only ESTs having a better top-hit bit score in the Ciliophora BLAST were retained for further analysis (any hits with e-values above 0.001 were considered insignificant).

#### OrthoVenn2

Candidate ciliate ESTs were also annotated in OrthoVenn2 (Xu et al., 2019) in order to identify orthologous clusters with *T. thermophila* genes (input sequences were translated in the ciliate code, and default settings were used). *Tetrahymena thermophila* orthologs identified in this step were classified as predicted ciliate ESTs if they also met the NCBI BLASTP selection criteria outlined above. A final subset of unique *T. thermophila* orthologs from each input transcriptome was generated by selecting the best match (based first on score and then on length) in any cases where multiple, redundant orthologs were returned. Pipeline-selected unique *T. thermophila* orthologs from each screened flatworm transcriptome, along with their GhostKOALA, NCBI BLASTP, and OrthoVenn2 scores, are listed in Table S2.

#### False positive filtering

The false-positive rate for the pipeline was estimated by screening transcriptomes from five reference eukaryotes considered highly unlikely to be ciliate hosts (see Supplementary Note 1). Only *T. thermophila* orthologs from flatworm EST libraries returning a significantly greater number of predicted ciliate ESTs than expected based upon the calculated false-positive rate (Table S3) were included in phylogenetic analyses.

### Characterization of predicted ciliate ESTs from flatworm transcriptomes

#### BLASTP analysis of predicted ciliate ESTs

In addition to the BLASTP step above, in which standard code translations were aligned against only the NCBI Platyhelminthes database, pipeline-selected unique *T. thermophila* orthologs were also aligned against the entire nonredundant protein database with Ciliophora sequences excluded (e-value threshold for significance set at 0.001). Bit scores were then compared for the top hit from this BLAST and the top hit from the corresponding BLAST with the ciliate code translation queried against only the Ciliophora database.

#### Stop codon analysis of predicted ciliate ESTs

The number of (in-frame) stop codons within the aligned regions of top Ciliophora-only BLASTP hits were compared for standard code and ciliate code translations of all pipeline-selected unique *T. thermophila* orthologs.

#### Kaiju classification of predicted ciliate ESTs

Independent taxonomic classification in Kaiju (Menzel et al., 2016) was completed for all pipeline-selected unique *T. thermophila* orthologs from enriched transcriptomes. This analysis was conducted in Greedy mode, which allows for mismatches in aligned protein sequences, using default parameters and NCBI BLAST *nr+euk* (non-redundant proteins for Archaea, bacteria, viruses, fungi, and microbial eukaryotes) as the reference database.

#### Genome BLASTs for predicted ciliate ESTs

Pipeline-selected unique *T. thermophila* orthologs from *S. mediterranea* transcriptomes (sexual and asexual) were aligned against both the *T. borealis* and *S. mediterranea* genomes by BLASTN (genome sequence files were retrieved from the Tetrahymena Genome Database (Stover et al., 2012) and PlanMine (Rozanski et al., 2019)). An equivalent number of randomly selected *S. mediterranea* sexual and asexual transcripts excluded at the GhostKoala step of the pipeline were analyzed as controls.

### Phylogenetic analyses

#### Phylogenetic analysis of predicted ciliate ESTs from flatworm transcriptomes

Pipeline-predicted ciliate ESTs from enriched flatworm transcriptomes were assigned orthology to genes from *Tetrahymena thermophila* and six other reference ciliates (*Ichthyophthirius multifiliis, Paramecium tetraurelia, Pseudocohnilembus persalinus, Tetrahymena borealis, Tetrahymena elliotti*, and *Tetrahymena malaccensis*) using OrthoVenn2. Predicted protein sequences within each orthogroup were then aligned by MAFFT (Katoh et al., 2002) using default parameters. Alignments were concatenated for computational efficiency to form 33 sets of 99 markers each, with a remaining set of 74 markers. A molecular phylogeny was inferred for each set of concatenated markers in IQ-TREE (Chernomor et al., 2016), using default parameters, a partition for each marker, and 10,000 bootstrap replicates. Finally, each of the resulting majority consensus tree topologies for the 34 alignments were prepared as a binary character data matrix in PAUP* 4.0 (Wilgenbusch and Swofford, 2003). Matrix representations were concatenated and analyzed to produce a matrix representation with likelihood (MRL) supertree. IQ-TREE was again utilized under default parameters, 10,000 bootstrap replicates, and with a data partition for each of the 34 subcomponents to the total binary character matrix. PAUP* 4.0 was used to generate the majority rule consensus supertree shown in Figure 1E.

#### cox1 phylogenetic analysis

Mitochondrial *cox1* sequences from cultured ciliates and reference species were aligned by MAFFT, using REVTRANS 2.0 (Wernersson and Pedersen, 2003) to ensure protein-coding information (translation table 4) was incorporated in the scaffold for multiple sequence alignment. IQ-TREE was next used to compute a maximum likelihood molecular phylogeny from the aligned nucleotide sequences, with 10,000 bootstrap replicates to determine branch support. The majority rule consensus topology in Figure 2 was prepared using PAUP* 4.0.

#### Platyhelminthes-Ciliophora cophylogeny

Individual flatworm and ciliate phylogenies were first prepared by converting NCBI Taxonomy data, results from prior evolutionary analyses of the Platyhelminthes (Álvarez-Presas et al., 2008; Laumer et al., 2015), and results from the ciliate phylogenies generated in the current study (Figure 1E and Figure 2) into partitioned, binary character matrices and computing relationships in IQ-TREE following the methods described above. Branches not supporting flatworm-ciliate associations were pruned, and PAUP*-generated trees were combined in TreeMap to produce the tanglegram shown in Figure 3. Jane 4.0 (Conow et al., 2010) was finally used to conduct an event-based codiversification analysis with random tip mapping, 100 generations, and a population size of 200. Results were analyzed in stats mode with 100 generations and a population size of 100 under a variety of different cost schemes (Table S4). Because the data did not conform to a normal distribution in all cases, the nonparametric Kruskal-Wallis test (with Bonferroni correction) was employed to determine statistical significance.

### *Tetrahymena* culture

#### Planarian sources

*Dugesia japonica* and *Schmidtea mediterranea* (asexual) were from clonal lines maintained in the laboratory for over 20 years. *Phagocata velata* was obtained from a vernal pool in New Hampshire (42°59’37.2”N 72°18’32.6”W). *Phagocata gracilis* and *Procotyla fluviatilis* were obtained from Ward’s Science (catalog numbers 470176-452 and 470180-238, respectively). *Girardia* sp. were obtained from Ward’s Science (catalog number 470176-558) and Carolina Biological Supply Company (catalog number 132954); these are thought to be *G. dorotocephala* and *G. tigrina*, respectively, but the species name is not specified because of the difficulty in distinguishing the two. *Bipalium adventitium* and *Bipalium kewense* were from laboratory populations descended from animals collected in New York in 2017 and Louisiana in 2018, respectively (both invasive distributions).

#### Mealworm cultures

Planarians were cut into small fragments in petri dishes filled with reverse osmosis-purified tap water adjusted to pH 7.0 and supplemented with a final concentration of 1.6 mM NaCl, 1 mM CaCl_2_, 1 mM MgSO_4_, 0.1 mM MgCl_2_, 0.1 mM KCl, and 1.2 mM NaHCO_3_ (‘planarian water’). Fragments were then exposed to a red LED (625 nm) for 24-48 hours at room temperature (for reasons that are unclear, we found light exposure encourages *Tetrahymena* to exit from planarian tissues). If ciliates were visible under a dissecting microscope, the flatworm fragments were removed, and a crushed, freeze-dried mealworm was added to the dish. 1 mL of the resulting culture was pelleted after 48 hours at 20ºC for DNA barcoding.

#### Neff cultures

Mealworm cultures of *Tetrahymena* (see above) were pelleted and resuspended at an initial density of 1,000 cells per 6 mL in Neff medium (Cassidy-Hanley et al., 1997), with a final concentration of 250 U/mL penicillin-streptomycin (Thermo-Fisher Scientific, catalog number 15140122). Growth at 20 ºC was monitored daily by placing a 1 μl drop of the culture on a glass slide, adding a drop of Protoslo (Carolina Biological, catalog number 885141), and counting the total number of cells under a compound microscope (at higher densities, cultures were diluted 1:10 prior to counting).

### cox1 DNA barcoding

Pelleted mealworm cultures were resuspended in 200 μl phosphate-buffered saline, and total DNA was purified with the Qiagen DNeasy Blood and Tissue kit (catalog number 69504), following the manufacturer’s protocol for extraction from crude lysates. Purified DNA was used as a template to amplify mitochondrial *cox1* by PCR. PCR products were purified using the Takara Bio NucleoSpin Gel and PCR Clean-Up kit (catalog number 740609) and sequenced using the forward PCR primer.

*cox1* forward primer: 5’ ATGTGAGTTGATTTTATAGA 3’

*cox1* reverse primer: 5’ CTCTTCTATGTCTTAAACCAGGCA 3’

### Paromomycin treatment

Animals were treated with a final concentration of 10 mM paromomycin in planarian water for 24 hours and then maintained in 1 mM paromomycin in planarian water for 28 days (solution was changed every four days).

### PCR assay for *Tetrahymena mobilis*

Single planarians were microwaved in 1 mL planarian water in microfuge tubes for 60 seconds and macerated with a glass rod. Macerates were then pelleted, resuspended in urea lysis buffer (350 mM NaCl, 10 mM Tris-HCl pH 7.4, 10 mM EDTA, 1% SDS, 42% urea), and phenol/chloroform extracted. Total DNA was precipitated with isopropyl alcohol, resuspended in TE buffer, and used as a template to amplify *S. mediterranea eRF1* and *T. mobilis HSP70* by PCR.

*eRF1* forward primer: 5’ CATTACCATCCCGGCTGATGAATATGGCACAGC 3’

*eRF1* reverse primer: 5’ CCAATTCTACCCGCCTTCTGCTGATTTGTCACTC 3’

*HSP70* forward primer: 5’ AGAACTCTGTCTCTCCCTCTTC 3’

*HSP70* reverse primer: 5’ ACCGTCGAGGTTGAACTTAC 3’

## Supporting information

Figure 2 Source Data (COX1 Sequences)

Figure 3 Source Data (Platyhelminthes Tree)

## ACKNOWLEDGEMENTS

This work was supported by New Hampshire-INBRE through an Institutional Development Award (P20GM103506) from the National Institute of General Medical Sciences of the National Institutes of Health. We thank Joseph Sevigny for bioinformatics support; Peter Ducey for providing *Bipalium* specimens; Andrei Rozanski and Jochen Rink for providing filtered EST data; Denis Lynn, F. Paul Doerder, and Aaron Aunins for technical advice; and Timothy Hastings, John Dustin, Callum Yule, Adam Willhoit, and Trisha Schuman for assistance with preliminary experiments.

## AUTHOR CONTRIBUTIONS

M.R.W. and J.P. conceived and designed the project. M.R.W., K.P., and K.S. performed the bioinformatic analyses and experiments. M.R.W. and J.P. interpreted the results. J.P. prepared the figures and wrote the manuscript.

## REFERENCES

Álvarez-Presas M, Baguñà J, Riutort M. 2008. Molecular phylogeny of land and freshwater planarians (Tricladida, Platyhelminthes): from freshwater to land and back. Molecular Phylogenetics and Evolution 47:555–568. doi:10.1016/j.ympev.2008.01.032

André E. 1909. Sur un nouvel Infusoire parasite des Dendrocoeles, Ophryoglena parasitica sp. n. Revue Suisse de Zoologie 17:273–280. doi:10.5962/bhl.part.3775

Armitage MJ, Young JO. 1990. A field and laboratory study of the parasites of the triclad Phagocata vitta (Duges). Freshwater Biology 24:101–107. doi:10.1111/j.1365-2427.1990.tb00311.x

Astrofsky KM, Schech JM, Sheppard BJ, Obenschain CA, Chin AM, Kacergis MC, Laver ER, Bartholomew JL, Fox JG. 2002. High mortality due to Tetrahymena sp. infection in laboratory-maintained zebrafish (Brachydanio rerio). Comparative Medicine 52:363–367.

Basson L, Van As J. 2006. Trichodinidae and other ciliophorans (phylum Ciliophora). Fish Diseases and Disorders, Volume 1: Protozoan and Metazoan Infections. CABI Books 154– 182. doi:10.1079/9780851990156.0154

Batson BS, Beale GH. 1983. Tetrahymena dimorpha sp.nov. (Hymenostomatida: Tetrahymenidae), a new ciliate parasite of Simuliidae (Diptera) with potential as a model for the study of ciliate morphogenesis. Philosophical Transactions of the Royal Society B: Biological Sciences 301:345–363. doi:10.1098/rstb.1983.0027

Batson BS, Rees FG. 1985. A paradigm for the study of insect-ciliate relationships: Tetrahymena sialidos sp. nov. (Hymenostomatida: Tetrahymenidae), parasite of larval Sialis Lutaria (Linn.) (Megaloptera: Sialidae). Philosophical Transactions of the Royal Society B: Biological Sciences 310:123–144. doi:10.1098/rstb.1985.0102

Bharti D, Kumar S, La Terza A, Chandra K. 2020. Dispersal of ciliated protist cysts: mutualism and phoresy on mites. Ecology 101:e03075. doi:10.1002/ecy.3075

Brandl MT, Rosenthal BM, Haxo AF, Berk SG. 2005. Enhanced survival of Salmonella enterica in vesicles released by a soilborne Tetrahymena species. Applied and Environmental Microbiology 71:1562–1569. doi:10.1128/AEM.71.3.1562-1569.2005

Brooks WM. 1968. Tetrahymenid ciliates as parasites of the gray garden slug. Hilgardia 39:205–276. doi:10.3733/hilg.v39n08p205

Cassidy-Hanley D, Bowen J, Lee JH, Cole E, VerPlank LA, Gaertig J, Gorovsky MA, Bruns PJ. 1997. Germline and somatic transformation of mating Tetrahymena thermophila by particle bombardment. Genetics 146:135–147. doi:10.1093/genetics/146.1.135

Chantangsi C, Lynn DH, Brandl MT, Cole JC, Hetrick N, Ikonomi P. 2007. Barcoding ciliates: a comprehensive study of 75 isolates of the genus Tetrahymena. International Journal of Systemic and Evolutionary Microbiology 57:2412–2423. doi:10.1099/ijs.0.64865-0

Chernomor O, von Haeseler A, Minh BQ. 2016. Terrace aware data structure for phylogenomic inference from supermatrices. Systematic Biology 65:997–1008. doi:10.1093/sysbio/syw037

Conow C, Fielder D, Ovadia Y, Libeskind-Hadas R. 2010. Jane: a new tool for the cophylogeny reconstruction problem. Algorithms for Molecular Biology 5:16. doi:10.1186/1748-7188-5-16

Corliss JO. 1970. The comparative systematics of species comprising the Hymenostome ciliate genus Tetrahymena. The Journal of Protozoology 17:198–209. doi:10.1111/j.1550-7408.1970.tb02356.x

Corliss JO. 1960. Tetrahymena chironomi sp.nov., a ciliate from midge larvae, and the current status of facultative parasitism in the genus Tetrahymena. Parasitology 50:111–153. doi:10.1017/S0031182000025245

Dickerson HW, Findly RC. 2014. Immunity to Ichthyophthirius infections in fish: a synopsis. Developmental and Comparative Immunology 43:290–299. doi:10.1016/j.dci.2013.06.004

Doerder FP. 2019. Barcodes reveal 48 new species of Tetrahymena, Dexiostoma, and Glaucoma: Phylogeny, ecology, and biogeography of new and established species. Journal of Eukaryotic Microbiology 66:182–208. doi:10.1111/jeu.12642

Elliott AM, Addison MA, Carey SE. 1962. Distribution of Tetrahymena pyriformis in Europe. The Journal of Protozoology 9:135–141. doi:10.1111/j.1550-7408.1962.tb02595.x

Ferguson H, Hicks B, Lynn D, Ostland V, Bailey J. 1987. Cranial ulceration in Atlantic salmon Salmo salar associated with Tetrahymena sp. Diseases of Aquatic Organisms 2:191–195. doi:10.3354/dao002191

Greider CW, Blackburn EH. 1985. Identification of a specific telomere terminal transferase activity in Tetrahymena extracts. Cell 43:405–413. doi:10.1016/0092-8674(85)90170-9

Grohme MA, Schloissnig S, Rozanski A, Pippel M, Young GR, Winkler S, Brandl H, Henry I, Dahl A, Powell S, Hiller M, Myers E, Rink JC. 2018. The genome of Schmidtea mediterranea and the evolution of core cellular mechanisms. Nature 554:56–61. doi:10.1038/nature25473

Haag WR, Williams JD. 2014. Biodiversity on the brink: an assessment of conservation strategies for North American freshwater mussels. Hydrobiologia 735:45–60. doi:10.1007/s10750-013-1524-7

He X, Lindsay-Mosher N, Li Y, Molinaro AM, Pellettieri J, Pearson BJ. 2017. FOX and ETS family transcription factors regulate the pigment cell lineage in planarians. Development 144:4540–4551. doi:10.1242/dev.156349

Hoffman GL, Landolt M, Camper JE, Coats DW, Stookey JL, Burek JD. 1975. A disease of freshwater fishes caused by Tetrahymena corlissi Thompson, 1955, and a key for identification of holotrich ciliates of freshwater fishes. The Journal of Parasitology 61:217–223. 10.2307/3278995

Holz PH, Portas T, Donahoe S, Crameri S, Rose K. 2015. Mortality in northern corroboree frog tadpoles (Pseudophryne pengilleyi) associated with Tetrahymena-like infection. Australian Veterinary Journal 93:295–297. doi:10.1111/avj.12337

Horowitz S, Gorovsky MA. 1985. An unusual genetic code in nuclear genes of Tetrahymena. Proceedings of the National Academy of Sciences USA 82:2452–2455. doi:10.1073/pnas.82.8.2452

Islas-Ortega AG, Aguilar-Aguilar R. 2014. Trichodina mutabilis (Protozoa: Ciliophora: Trichodinidae) from the characid fish Astyanax mexicanus in the Cuatro Ciénegas region, northern Mexico. Revista Mexicana de Biodiversidad 85:613–616. doi:10.7550/rmb.33107

Ivankovic M, Haneckova R, Thommen A, Grohme MA, Vila-Farré M, Werner S, Rink JC. 2019. Model systems for regeneration: planarians. Development 146:dev167684. doi:10.1242/dev.167684

Jerome CA, Lynn DH, Simon EM. 1996. Description of Tetrahymena empidokyrea n.sp., a new species in the Tetrahymena pyriformis sibling species complex (Ciliophora, Oligohymenophorea), and an assessment of its phylogenetic position using small-subunit rRNA sequences. Canadian Journal of Zoology 74:1898–1906. doi:10.1139/z96-214

Justine J-L, Théry T, Gey D, Winsor L. 2019. First record of the invasive land flatworm Bipalium adventitium (Platyhelminthes, Geoplanidae) in Canada. Zootaxa 4656:zootaxa.4656.3.13. doi:10.11646/zootaxa.4656.3.13

Justine J-L, Winsor L, Gey D, Gros P, Thévenot J. 2018. Giant worms chez moi! Hammerhead flatworms (Platyhelminthes, Geoplanidae, Bipalium spp., Diversibipalium spp.) in metropolitan France and overseas French territories. PeerJ 6:e4672. doi:10.7717/peerj.4672

Kanana Y, Riutort M. 2019. First record of freshwater planarian Girardia tigrina (Platyhelminthes, Tricladida, Continenticola) in Eastern Europe. Zootaxa 4624:zootaxa.4624.4.13. doi:10.11646/zootaxa.4624.4.13

Kanehisa M, Sato Y, Morishima K. 2016. BlastKOALA and GhostKOALA: KEGG tools for functional characterization of genome and metagenome sequences. Journal of Molecular Biology 428:726–731. doi:10.1016/j.jmb.2015.11.006

Katoh K, Misawa K, Kuma K, Miyata T. 2002. MAFFT: a novel method for rapid multiple sequence alignment based on fast Fourier transform. Nucleic Acids Research 30:3059–3066. doi:10.1093/nar/gkf436

Kawakatsu M, Mitchell R, Hirao Y, Tanaka I. 1984. Occurrence of Dugesia dorotocephala (Woodworth, 1897)(Turbellaria, Tricladida, Paludicola) in Honolulu, Hawaii. Biological Magazine of Okinawa 22:1–9.

Knight DR, McDougle HC. 1944. A protozoan of the genus Tetrahymena found in the domestic fowl. American Journal of Veterinary Research 5:113–116.

Kruger K, Grabowski PJ, Zaug AJ, Sands J, Gottschling DE, Cech TR. 1982. Self-splicing RNA: autoexcision and autocyclization of the ribosomal RNA intervening sequence of Tetrahymena. Cell 31:147–157. doi:10.1016/0092-8674(82)90414-7

Laumer CE, Hejnol A, Giribet G. 2015. Nuclear genomic signals of the ‘microturbellarian’ roots of platyhelminth evolutionary innovation. eLife 4:e05503. doi:10.7554/eLife.05503

Leibowitz MP, Zilberg D. 2009. Tetrahymena sp. infection in guppies, Poecilia reticulata Peters:parasite characterization and pathology of infected fish. Journal of Fish Diseases 32:845–855. doi:10.1111/j.1365-2761.2009.01062.x

Lynn D, Doerder F, Gillis P, Prosser R. 2018. Tetrahymena glochidiophila n. sp., a new species of Tetrahymena (Ciliophora) that causes mortality to glochidia larvae of freshwater mussels (Bivalvia). Diseases of Aquatic Organisms 127:125–136. doi:10.3354/dao03188

Lynn D, Gransden SG, Wright A, Josephson G. 2000. Characterization of a new species of the ciliate Tetrahymena [Ciliophora: Oligohymenophorea] isolated from the urine of a dog: first report of Tetrahymena from a mammal. Acta Protozoologica 39:289–294.

Lynn DH, Molloy D, LeBrun R. 1981. Tetrahymena rotunda n. sp. (Hymenostomatida: Tetrahymenidae), a ciliate parasite of the hemolymph of Simulium (Diptera: Simuliidae). Transactions of the American Microscopical Society 100:134–141. doi:10.2307/3225796

McGurk ES, Ford S, Bushek D. 2016. Unusually abundant and large ciliate xenomas in oysters, Crassostrea virginica, from Great Bay, New Hampshire, USA. Journal of Invertebrate Pathology 137:23–32. doi:10.1016/j.jip.2016.04.001

Menzel P, Ng KL, Krogh A. 2016. Fast and sensitive taxonomic classification for metagenomics with Kaiju. Nature Communications 7:11257. doi:10.1038/ncomms11257

Merryman MS, Sánchez Alvarado A, Jenkin JC. 2018. Culturing planarians in the laboratory. Methods in Molecular Biology. Clifton, NJ 1774:241–258. doi:10.1007/978-1-4939-7802-1_5

Nanney DL, Simon EM. 2000. Laboratory and evolutionary history of Tetrahymena thermophila. Methods in Cell Biology 62:3–25. doi:10.1016/s0091-679x(08)61527-7

Örsted AS. 1844. Entwurf einer systematischen Einteilung und speciellen Beschreibung der Plattwuermer auf microscopische Untersuchungen gegründet J.C. Scharling, Copenhagen. 96 p.

Palmer E, Wilhelm JM. 1978. Mistranslation in a eucaryotic organism. Cell 13:329–334. doi:10.1016/0092-8674(78)90201-5

Petz W, Valbonesi A, Schiftner U, Quesada A, Cynan Ellis-Evans J. 2007. Ciliate biogeography in Antarctic and Arctic freshwater ecosystems: endemism or global distribution of species? FEMS Microbiology Ecology 59:396–408. doi:10.1111/j.1574-6941.2006.00259.x

Pinheiro MDO, Bols NC. 2014. Delineating cellular interactions between ciliates and fish by coculturing Tetrahymena thermophila with fish cells. Cell Biology International 38:1138–1147. doi:10.1002/cbin.10310

Pitsch G, Adamec L, Dirren S, Nitsche F, Šimek K, Sirová D, Posch T. 2017. The greenTetrahymena Utriculariae n. sp. (Ciliophora, Oligohymenophorea) with Its endosymbiotic algae (Micractinium sp.), living in traps of a carnivorous aquatic plant. Journal of Eukaryotic Microbiology 64:322–335. doi:10.1111/jeu.12369

Pokharel P, Ghimire R, Lamichhane P. 2021. Efficacy and safety of paromomycin for visceral leishmaniasis: A systematic review. Journal of Tropical Medicine 2021:8629039. doi:10.1155/2021/8629039

Preer JR, Preer LB, Rudman BM, Barnett AJ. 1985. Deviation from the universal code shown by the gene for surface protein 51A in Paramecium. Nature 314:188–190. doi:10.1038/314188a0

Prosser RS, Lynn DH, Salerno J, Bennett J, Gillis PL. 2018. The facultatively parasitic ciliated protozoan, Tetrahymena glochidiophila (Lynn, 2018), causes a reduction in viability of freshwater mussel glochidia. Journal of Invertebrate Pathology 157:25–31. doi:10.1016/j.jip.2018.07.012

Rahaman A, Miao W, Turkewitz AP. 2009. Independent transport and sorting of functionally distinct protein families in Tetrahymena thermophila dense core secretory granules. Eukaryotic Cell 8:1575–1583. doi:10.1128/EC.00151-09

Rataj M, Vd’ačný P. 2021. Cryptic host-driven speciation of mobilid ciliates epibiotic on freshwater planarians. Molecular Phylogenetics and Evolution 161:107174. doi:10.1016/j.ympev.2021.107174

Rataj M, Vďačný P. 2020. Multi-gene phylogeny of Tetrahymena refreshed with three new histophagous species invading freshwater planarians. Parasitology Research 119:1523–1545. doi:10.1007/s00436-020-06628-0

Rataj M, Vd’ačný P. 2019. Living morphology and molecular phylogeny of oligohymenophorean ciliates associated with freshwater turbellarians. Diseases of Aquatic Organisms 134:147–166. doi:10.3354/dao03366

Reynoldson TB. 1956. The population dynamics of host specificity in Urceolaria mitra (Peritricha) epizoic on fresh-water triclads. Journal of Animal Ecology 25:127–143. doi:10.2307/1854

Rozanski A, Moon H, Brandl H, Martín-Durán JM, Grohme MA, Hüttner K, Bartscherer K, Henry I, Rink JC. 2019. PlanMine 3.0—improvements to a mineable resource of flatworm biology and biodiversity. Nucleic Acids Research 47:D812–D820. doi:10.1093/nar/gky1070

Sharon G, Pimenta Leibowitz M, Chettri JK, Isakov N, Zilberg D. 2014. Comparative study of infection with Tetrahymena of different ornamental fish species. Journal of Compartive Pathology 150:316–324. doi:10.1016/j.jcpa.2013.08.005

Shaw S, Speare R, Lynn DH, Yeates G, Zhao Z, Berger L, Jakob-Hoff R. 2011. Nematode and ciliate nasal infection in captive archey’s frogs (Leiopelma archeyi). Journal of Zoo and Wildlife Medicine 42:473–479. doi:10.1638/2010-0180.1

Shumway W. 1940. A ciliate protozoon parasitic in the central nervous system of larval ambystoma. Biological Bulletin 78:283–288. doi:10.2307/1537778

Sluys R, Kawakatsu M, Yamamoto K. 2010. Exotic freshwater planarians currently known from Japan. Belgian Journal of Zoology 140:103–109.

Souidenne D, Florent I, Dellinger M, Justine JL, Romdhane MS, Furuya H, Grellier P. 2016. Diversity of apostome ciliates, Chromidina spp. (Oligohymenophorea, Opalinopsidae), parasites of cephalopods of the Mediterranean Sea. Parasite 23:33. doi:10.1051/parasite/2016033

Stout JD. 1954. The ecology, life history and parasitism of Tetrahymena [Paraglaucoma] rostrata (Kahl) Corliss. The Journal of Protozoology 1:211–215. doi:10.1111/j.1550-7408.1954.tb00819.x

Stover NA, Punia RS, Bowen MS, Dolins SB, Clark TG. 2012. Tetrahymena genome database Wiki: a community-maintained model organism database. Database 2012:bas007. doi:10.1093/database/bas007

Strüder-Kypke MC, Wright A-DG, Jerome CA, Lynn DH. 2001. Parallel evolution of histophagy in ciliates of the genus Tetrahymena. BMC Evolutionary Biology 1:5. doi:10.1186/1471-2148-1-5

Swapna LS, Molinaro AM, Lindsay-Mosher N, Pearson BJ, Parkinson J. 2018. Comparative transcriptomic analyses and single-cell RNA sequencing of the freshwater planarian Schmidtea mediterranea identify major cell types and pathway conservation. Genome Biology 19:124. doi:10.1186/s13059-018-1498-x

The Galaxy Community. 2022. The Galaxy platform for accessible, reproducible and collaborative biomedical analyses: 2022 update. Nucleic Acids Research 50:W345–W351. doi:10.1093/nar/gkac247

Ushida K. 2018. Symbiotic methanogens and rumen ciliates In: Hackstein JHP, editor. (Endo)Symbiotic Methanogenic Archaea, Microbiology Monographs. Cham: Springer International Publishing. pp. 25–35. doi:10.1007/978-3-319-98836-8_3

Wernersson R. 2006. Virtual Ribosome--a comprehensive DNA translation tool with support for integration of sequence feature annotation. Nucleic Acids Research 34:W385–388. doi:10.1093/nar/gkl252

Wernersson R, Pedersen AG. 2003. RevTrans: multiple alignment of coding DNA from aligned amino acid sequences. Nucleic Acids Research 31:3537–3539. doi:10.1093/nar/gkg609

Wilgenbusch JC, Swofford D. 2003. Inferring evolutionary trees with PAUP*. Current Protocols in Bioinformatics 00:6.4.1-6.4.28. doi:10.1002/0471250953.bi0604s00

Wright JF. 1981. Tetrahymena pyriformis (Ehrenberg) and T. corlissi Thompson parasitic in stream-dwelling triclads (Platyhelminthes: Turbellaria). The Journal of Parasitology 67:131–133. doi:10.2307/3280799

Wright JF. 1969. The ecology of stream-dwelling triclads. Dissertation University of Wales.

Xu L, Dong Z, Fang L, Luo Y, Wei Z, Guo H, Zhang G, Gu YQ, Coleman-Derr D, Xia Q, Wang Y. 2019. OrthoVenn2: a web server for whole-genome comparison and annotation of orthologous clusters across multiple species. Nucleic Acids Research 47:W52–W58. doi:10.1093/nar/gkz333

Yang W, Jiang C, Zhu Y, Chen K, Wang G, Yuan D, Miao W, Xiong J. 2019. Tetrahymena Comparative Genomics Database (TCGD): a community resource for Tetrahymena. Database 2019:baz029. doi:10.1093/database/baz029

