## Supplementary Information for "Flatworm transcriptomes reveal widespread parasitism by histophagous ciliates"

### SUPPLEMENTARY NOTES

#### Supplementary Note 1: False positive rate calculation for bioinformatic pipeline

We applied two selection criteria in choosing the five reference species used to estimate the false-positive rate for our bioinformatic pipeline (*Amphidinium carterae, Euglena gracilis, Physarum polycephalum, Sargassum vulgare,* and *Ulva lactuca*). First, like all of the screened Platyhelminthes species, they had to be eukaryotes that were excluded from the KEGG database, because we found this impacted the initial identification of candidate ciliate ESTs using GhostKOALA. Second, they had to be unlikely ciliate hosts. The broad distribution of known Ciliophora endosymbionts, including within animals inhabiting brackish and marine environments (McGurk et al., 2016; Souidenne et al., 2016), makes it is difficult to place any organism into this category with complete confidence. Nonetheless, we are unaware of any documented ciliate endosymbionts in unicellular (*A. carterae* and *E. gracilis*) or acellular (*P. polycephalum*) eukaryotes, or in marine algae (*S. vulgare* and *U. lactuca*). The calculated false-positive rate obtained from these reference transcriptomes (eight out of 10,000 ESTs) set, if anything, an overly stringent threshold, as 15 of the screened Platyhelminthes libraries were significantly *depleted* in predicted ciliate ESTs in comparison (Table S3, pink). We accepted the resulting potential for false-negatives in deducing infected flatworm species because we were more concerned with the possibility that inclusion of false-positive hosts would produce misleading phylogenetic reconstructions. The significant differences between predicted ciliate transcripts from enriched and non-enriched libraries (Figure 1B,C) also suggests our false-positive threshold was effective in distinguishing infected and uninfected species.

#### Supplementary Note 2: Impact of transcriptome assembly methods on pipeline results

Lack of transcriptome enrichment in predicted ciliate ESTs (Table 1) should not be construed as evidence that a particular flatworm species (or population) is uninfected, for a variety of reasons. Most importantly, the transcriptomes of multicellular organisms are routinely filtered to remove contaminating sequences from resident microbiota. Depending on the parameters involved, this could include unicellular ciliates. As a preliminary assessment of how this might have impacted our results, we obtained filtered ESTs from seven of the screened Platyhelminthes libraries [*D. lacteum, D. japonica, P. torva, P. nigra, P. tenuis, S. mediterranea* (sexual), and *S. polychroa*] and ran them through our pipeline. This did yield a significant increase in the number of predicted ciliate ESTs for two of the three already enriched transcriptomes (*D. lacteum* and *P. torva* – the additional recovered sequences were *not* included in further analyses). However, it failed to put any of the non-enriched transcriptomes above the threshold required to exclude a false-positive. It is also noteworthy that 750 putative *Tetrahymena* transcripts were removed from the screened *S. mediterranea* (asexual) transcriptome prior to our pipeline analysis (Swapna et al., 2018), yet we were still able to identify an additional 754 predicted ciliate ESTs. Given our high confidence in the recovery of bona fide ciliate transcripts from enriched flatworm species (Results), these observations indicate commonly employed transcriptome assembly methods do not remove all Ciliophora sequences, likely in part because they fail to account for the nonstandard ciliate genetic code. We cannot, however, exclude the possibility that some of the remaining non-enriched libraries from our study constitute false-negatives due to filtering of input ESTs.

**Supplementary Note 3: Phylogenetic analysis of pipeline-predicted ciliate ESTs**

One possible complicating factor of the phylogenetic analysis of pipeline-derived ESTs (and any future research incorporating this approach) is the potential for concurrent infection of a given host by two or more ciliate species [e.g., four associations have been described for *D. lacteum* (André, 1909; Örsted, 1844; Reynoldson, 1956; Wright, 1969)]. In any such situation, it would be difficult to assign predicted Ciliophora ESTs to particular species when concatemerizing transcripts (this would apply whether multiple ciliate species were present in a given animal, or different animals from a population used to generate a transcriptome were infected by different ciliates). The high bootstrap values we obtained mitigate this concern to some degree, but we cannot rule these possibilities out entirely based on existing data. Regardless, our central conclusion that ciliates infect additional flatworm species would not be affected.

### SUPPLEMENTARY TABLE LEGENDS

**Table S1: Screened flatworm transcriptomes.** Input transcriptomes for the pipeline analysis and corresponding numbers of ESTs.

**Table S2: Unique *T. thermophila* orthologs from flatworm transcriptomes.** Pipeline-identified, unique *T. thermophila* orthologs are listed for flatworm species enriched (top) or not enriched (bottom, orange) in predicted ciliate ESTs. NCBI BLASTP scores are for the ciliate translation against the Ciliophora database. Sequences meeting criteria for predicted ciliate ESTs in multiple reading frames (eight) are listed separately here but are counted only once in Figure 1, Table 1, and Table S3.

**Table S3: Flatworm transcriptomes enriched in predicted ciliate ESTs.** Numbers of ESTs recovered at each step of the pipeline are indicated for all screened species. Those significantly enriched or not enriched in predicted ciliate ESTs, relative to the combined dataset for the five reference species, are shaded blue and pink, respectively (see Supplementary Note 1 for details on selection of reference species). Statistical significance was determined using 2x2 chi-square tests, with a Bonferroni corrected p-value threshold of 1.47E-03 (i.e., 0.05/34 transcriptomes). Yates' correction (Y) was applied when any expected frequency was <10.

**Table S4: Statistical analysis of coevolution events.** Statistical significance was determined for cophylogenies reconstructed in Jane under the indicated cost schemes using the Kruskal-Wallis test with Bonferroni correction. The differences between randomized costs (RTM = random tip mapping; RPT = random parasite trees) and calculated global costs were significant for all schemes.
