## Supplementary material for "Flatworm transcriptomes reveal widespread parasitism by histophagous ciliates": Figures S1 and S2

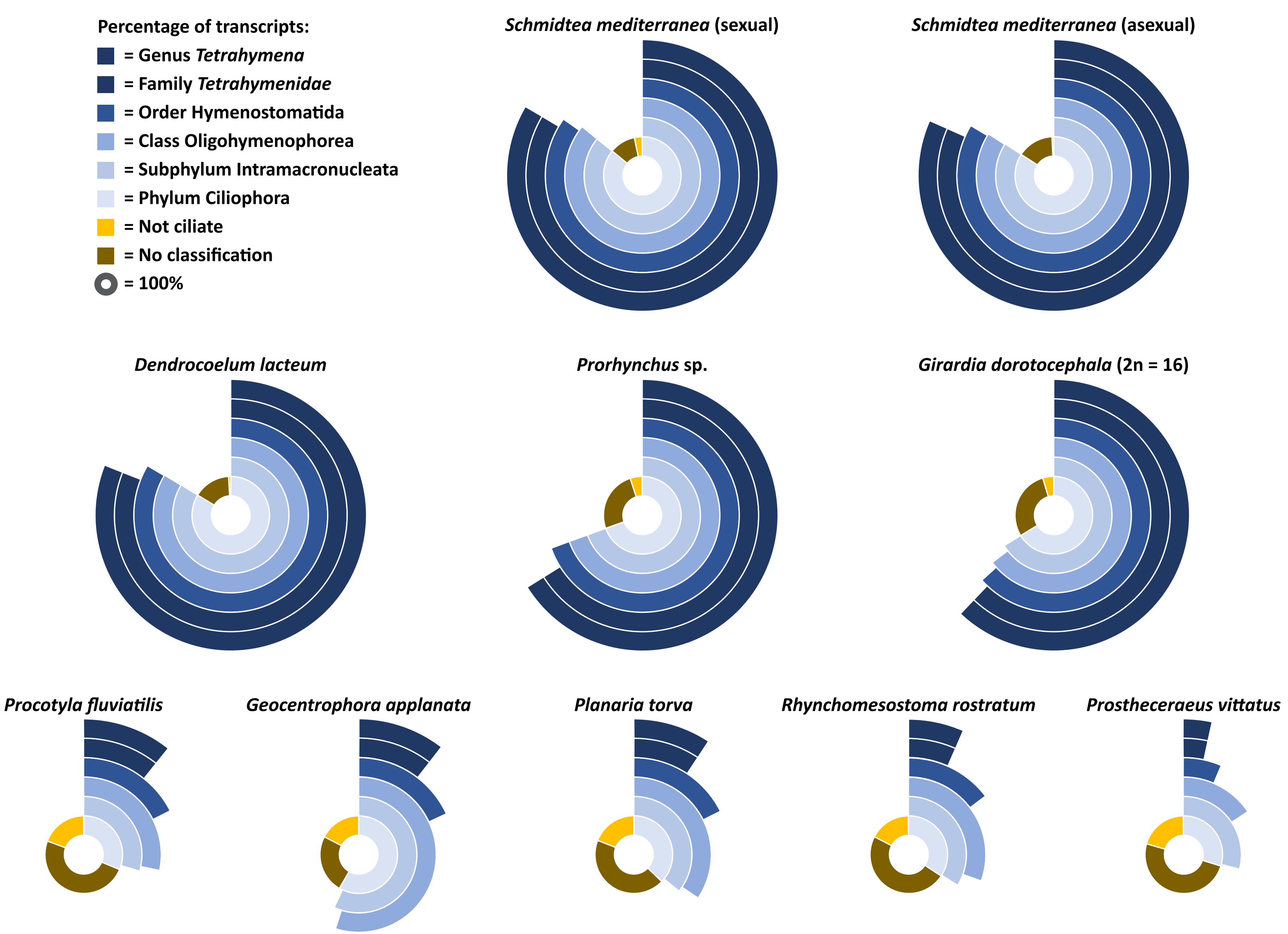


**Figure S1: Kaiju classification of ciliate ESTs from flatworm transcriptomes.** For every pipeline-screened transcriptome enriched in predicted ciliate ESTs (see Table 1), taxonomic classification of corresponding unique *T. thermophila* orthologs was conducted in Kaiju, with NCBI nonredundant proteins for Archaea, bacteria, viruses, fungi, and microbial eukaryotes as the reference database. Mismatches in aligned protein sequences were allowed. Sunburst plots show the percentage of transcripts classified as indicated in the key for each flatworm species (numbers are included in the source data file).

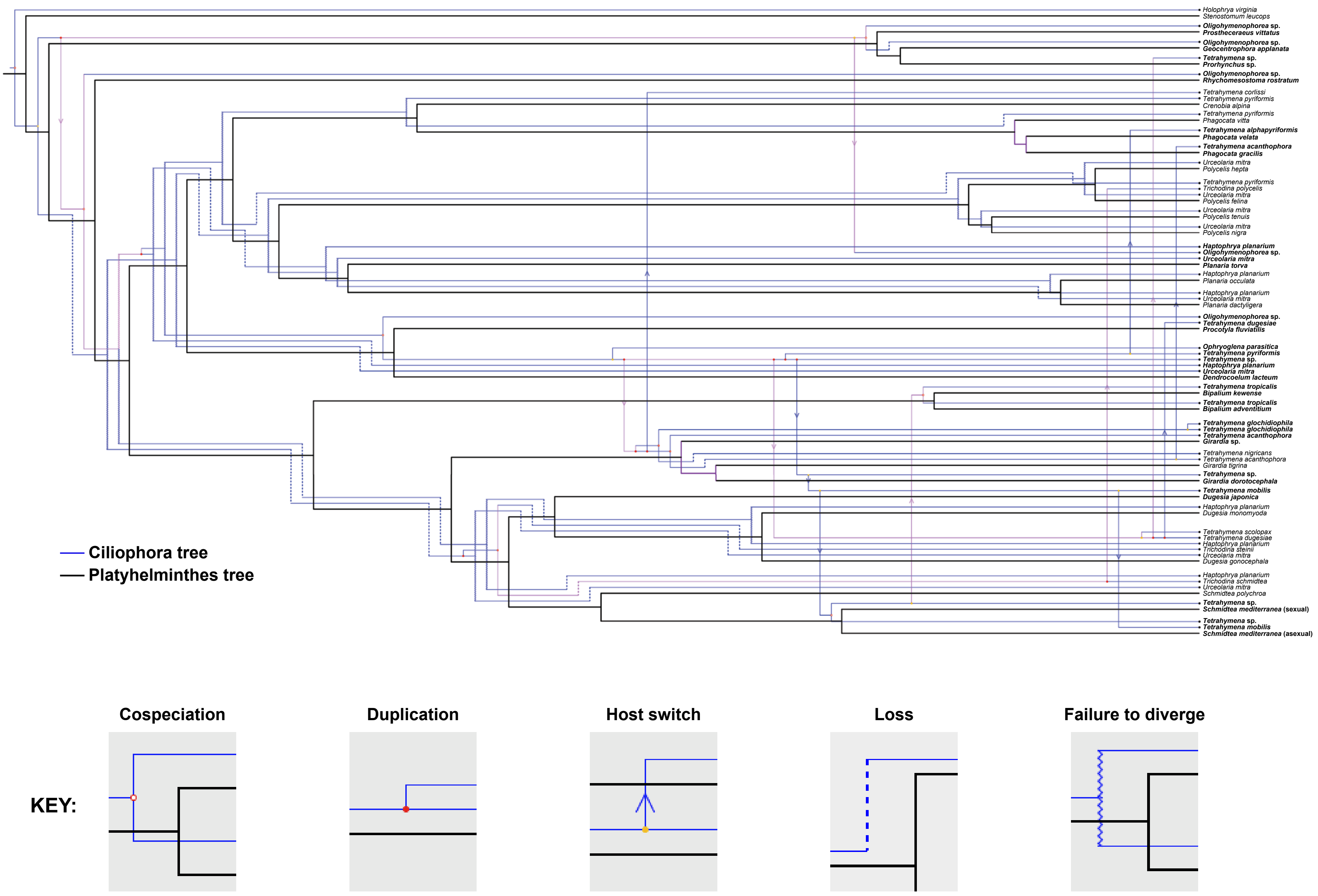


**Figure S2: Phylogenic reconciliation for ciliate parasites and their flatworm hosts.** Most frequently recovered cophylogenetic topology generated by Jane (150 of 917 equal-cost solutions), using default parameters (100 generations, population size of 200, and Jane default cost setting). Bold species denote associations documented in this work. See Table S4 for summary of statistical analyses.
