## Supplementary figures and images for "Flatworm transcriptomes reveal widespread parasitism by histophagous ciliates"

### Figure 4C Source Data

**Untreated**

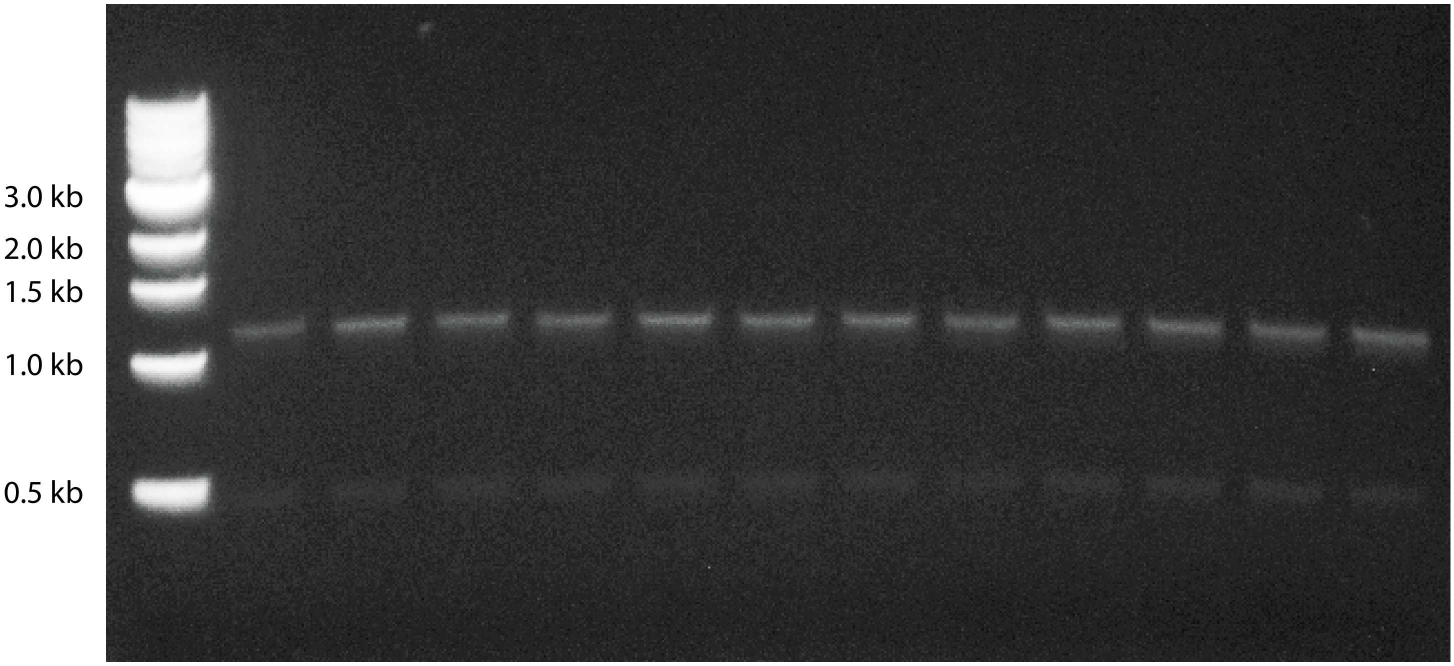


**Paromomycin-treated**

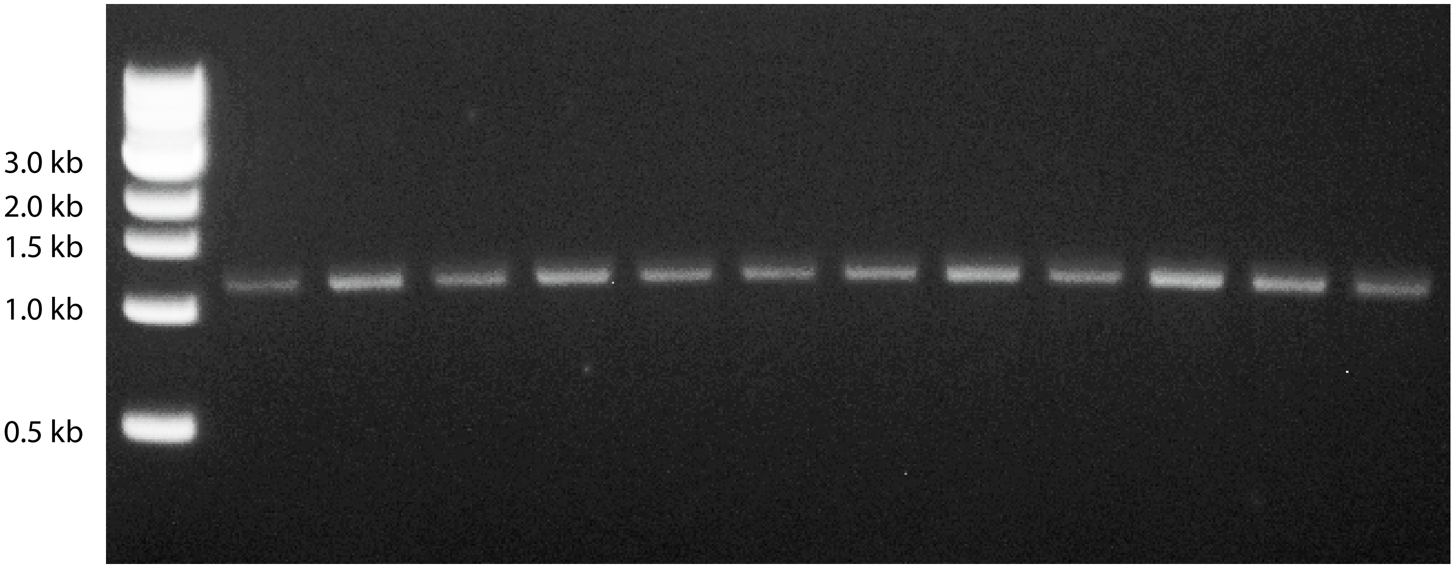
